# Transcription-factor binding to replicated DNA

**DOI:** 10.1101/2019.12.15.877035

**Authors:** Raz Bar-Ziv, Sagie Brodsky, Michal Chapal, Naama Barkai

## Abstract

Genome replication perturbs the DNA regulatory environment by displacing DNA-bound proteins, replacing nucleosomes, and introducing dosage-imbalance between regions replicating at different S phase stages. Recently, we showed that these effects are integrated to maintain transcription homeostasis: replicated genes increase in dosage, but their expression remains stable due to replication-dependent epigenetic changes that suppress transcription. Here, we examined whether reduced transcription from replicated DNA results from limited accessibility to regulatory factors, by measuring the time-resolved binding of RNA polymerase II (RNAPII) and specific transcription factors (TFs) to DNA during S phase in budding yeast. We show that RNAPII binding-pattern is largely insensitive to DNA dosage, indicating limited binding to replicated DNA. By contrast, binding of three TFs (Reb1, Abf1 and Rap1) to DNA increased with the increasing DNA dosage. We conclude that the replication-specific chromatin environment remains accessible to regulatory factors, but suppresses RNA polymerase recruitment.

## Introduction

DNA serves as a common template connecting gene transcription and genome replication, two fundamental cellular processes. Transcription and replication both depend on the ability of regulatory factors to access the DNA and progress smoothly along the genome. Accessibility of DNA to regulatory factors is limited by its wrapping around histone octamers which form nucleosomes, the basic building blocks of chromatin. DNA accessibility is not uniform across the genome but depends on multiple elements, including the DNA sequence, regulatory factors present at adjacent regions, and modifications added to histone tails, all of which could affect the affinity of DNA to histones. Inevitably, this makes the chromatin environment a major effector of transcription and replication, explaining the extensive regulation of this environment during both processes [1, 2].

DNA replication challenges the stability of gene expression at several levels. First, replication introduces gene dosage imbalance, simply because replicated genes have double DNA dosage relative to those yet to replicate. Second, encounters between the progressing replication fork and DNA-bound transcription factors (TFs), or transcribing RNA polymerases, may impinge on both processes. Furthermore, replication perturbs the epigenetic landscape by directly modifying histones, and by introducing new histones that carry a unique, position-independent set of marks [3]. Some marks recover their pre-replication pattern rapidly, but others recover slowly, over a period that can extend beyond S-phase [4, 5]. Finally, nucleosome positions are also altered as histones are evicted by the progressing replication fork, and regain their pre-replication distribution with some delay [6–8]. Therefore, during DNA replication, gene transcription proceeds in a unique DNA environment that is different from the environment it encounters outside of S phase.

In bacteria, the gene-dosage imbalance introduced during DNA replication results in increased transcription of replicated genes, an effect incorporated into bacterial gene-regulatory strategies [9]. Eukaryotes, by contrast, buffer this dosage imbalance by suppressing transcription from replicated DNA [10–12]. We recently showed that in budding yeast, this buffering depends on the acetylation of H3K56 by the acetyltransferase Rtt109 [13], and on H3K4 methylation by the Paf1-recruited COMPASS complex [14]. Specifically, the histone mark H3K56ac is deposited in replicated regions [15, 16] and H3K4me3, which is associated with active transcription, decreases in replicated regions [14]. Notably, in the case of replication stress, this feedback is stabilized by the DNA replication checkpoint, underlining its functional relevance. Therefore, while the increased dosage of replicated genes can potentially lead to increased transcription, this is suppressed by the unique epigenetic landscape characterizing replicated DNA. The suppression of transcription from replicated DNA could result from limited ability of regulatory factors to access the DNA at these regions. Alternatively, it could result from a more specific inhibition of the transcription machinery itself. In this study, we set to distinguish between these possibilities by examining the unperturbed replication dynamics of the two intermediate layers connecting the chromatin environment to gene expression: binding of RNA polymerase II (RNAPII) to replicated DNA and binding of transcription factors (TFs) to their cis-regulatory elements in replicated gene promoters.

We followed budding yeast as they progressed synchronously through unperturbed S phase, profiling, at high temporal resolution, the genome-wide binding of RNAPII and of three TFs that have a large number of binding targets: Reb1, Abf1 and Rap1. We found little evidence for a replication-associated depletion of RNAPII from DNA, suggesting that, if evicted by the progressing fork, it resumes binding rapidly. Further, the increase in DNA content during replication had only a minor effect (~20%) on RNAPII binding to replicated genes or promoters. In contrast, the transient depletion of the three TFs examined from replicating regions was more pronounced. These factors then regained binding, increasing in abundance with the increasing DNA content of replicated regions to an extent that was at least proportional to the increase in DNA dosage. Together, our data suggests that the unique chromatin environment introduced during replication does not prevent TF binding but does suppress the ability of these factors to recruit RNAPII and initiate gene transcription.

## Results

### The pattern of RNAPII binding to DNA during S phase

RNAPII binds to gene promoters and subsequently elongates along the coding sequence. Replication can perturb the genomic distribution of RNAPII binding in at least two ways. First, RNAPII could be transiently evicted by the replication fork or associated factors. Second, RNAPII binding may increase in replicated regions, whose DNA dosage increases, at the expense of other regions yet to replicate.

To examine the extent to which DNA replication perturbs RNAPII binding to DNA, we analyzed our previously published data of RNAPII binding in cells progressing synchronously through S phase [5]. In this experiment, we released budding yeast from a-factor induced arrest at the end of G1, and sampled the culture at three-minute intervals for DNA content and RNAPII binding, using DNA-Seq and ChIP-Seq, respectively. We verified the synchronous progression of the cells through S-phase by DNA staining, and observed the expected correlation between RNAPII occupancy and absolute expression levels, as well as the rapid decrease in its binding to mating genes, whose expression is down-regulated upon release from arrest, and the sequential increase in binding to cell-cycle induced genes (Figure 1A&B).

**Figure 1:**
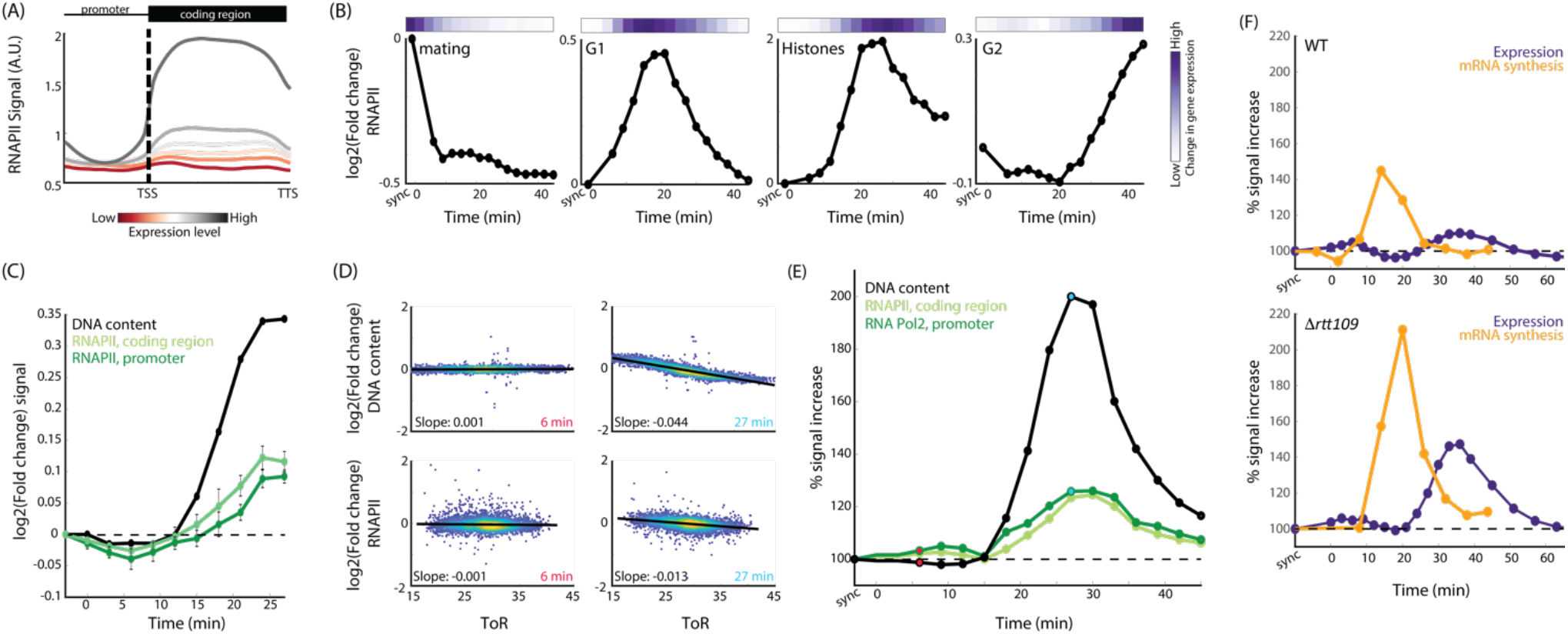
Genomic localization of RNAPII during DNA replication. (A) *RNAPII binding correlates with gene expression:* Genes were classified into six groups based on their absolute expression level, and were aligned by their transcription start and termination sites (TSS and TTS, respectively), and binned to control for variation in gene length (See methods). (TSS and TTS annotations as defined in David et al., 2006). Shown is the binding profile of RNAPII along the genes, averaged over each group. See also methods and Figure S1A. (B) *Temporal changes in RNAPII correlate with changes in mRNA expression:* The average (log2) fold change in the binding intensity of RNAPII to ORFs of the indicated gene groups is shown at different time points following release from G1 arrest (bottom). The corresponding expression change of the same gene group, as quantified in Voichek et al., 2016, is shown as color-code (top). See also Figure S1B for specific genes. (C) *RNAPII binding to early replicating genes:* RNAPII binding intensities and DNA abundance were log-normalized by their respective values in G1-arrested cells and averaged over the 200 earliest replicating genes. Shown are the respective averages, and the standard error, at subsequent time points following release from G1 arrest. (D) *Temporal progression of DNA replication:* Individual panels depict the measured fold change in gene DNA content (top) or RNAPII binding (bottom), normalized by the G1 arrest time points (synchronized), as a function of the respective ToR. Each dot represents a gene, and each plot corresponds to a specific time point following release from G1 arrest. Note the uniform behavior at early times, before replication starts, compared to later times, when cells have replicated part of their genome. This dependency between the measured parameter (DNA content or RNAPII binding) and ToR is quantified by the respective slopes, as shown. ToR of different genomic regions as defined by Yabuki et al., 2002 (see Methods). (E) Minor sensitivity of *RNAPII to gene dosage:* Shown is the time-dependent dependency of RNAPII binding on ToR, calculated on all genes with an annotated TSS as defined previously [28], quantified as in (D) and normalized by the corresponding dependency of DNA content in mid-S (see Methods, and Figure S1C). The red and blue dots correspond to the time points shown in (D). (F) *A complete loss of transcription buffering in RTT109-deleted cells:* Shown is the change in mRNA expression and mRNA synthesis rate, calculated as described in (D)&(E), for wild-type (top) and *□rtt109* (bottom) strains. mRNA synthesis rate was calculated as previously described in Voichek et al., 2016. Note that the plots represent separately-executed time course experiments. As each time course involves release from cell cycle synchronization, the time points between (E) and (F) are not directly comparable, due to slight temporal shifts.

To examine whether the progressing replication fork coincides with eviction of RNAPII from regions being replicated, we considered early-replicating regions, as defined by available data quantifying the time at S phase at which each genomic region is replicated (time of replication, ToR) [17]. Quantifying DNA content in these regions, we observed the expected increase shortly after the release from a-factor arrest (Figure 1C). By contrast, RNAPII binding was largely stable, showing only a minor transient reduction in binding to these regions. Therefore, within our temporal resolution (~3 minutes), we could not detect a significant depletion of RNAPII that is associated with the passing of the replication fork, suggesting that, if evicted, it regains binding rapidly.

Next, we asked if, following the passing of the replication fork, the increase in DNA content results in increased RNAPII binding. To this end, we needed to first quantify the progression of replication at each time point. We did this by examining the correlation between DNA content at each genomic region and the replication time (ToR) of this region. At early (G1) or late (G2/M) time points, the DNA content was uniform, independent of ToR. By contrast, at intermediate time points, DNA content increased in proportion to ToR (Figure 1D, top panel, and Figure S1C). The extent to which DNA content increases with ToR therefore defines the replication-dependent dosage bias (Figure 1E) and, accordingly, the expected increase in RNAPII binding: if no buffering occurs, and RNAPII increases precisely in proportion to DNA content, its dependency on ToR will mimic that of the DNA content. If, on the other hand, its binding is buffered against the dosage changes, it will remain independent of ToR throughout the time course. Indeed, applying this approach to re-analyze data of mRNA synthesis rates in wild-type cells, and in *Δrtt109* mutants that lose transcription buffering [13], captures the proportionality between transcription rates and DNA dosage in the *Δrtt109* mutant, and the minor dependency of mRNA synthesis rate on DNA dosage in wild-type cells (Figure 1F).

Quantifying the dependency of RNAPII binding on DNA dosage (Figure 1E), we find that RNAPII binding to replicated DNA increased by only ~20% relative to the increase in DNA content, similar to the change in mRNA synthesis observed in wild type cells (Figure 1E&F). Therefore, during unperturbed S phase, limited RNAPII binding to replicated DNA fully accounts for the buffering of transcription rates. Note that this limited increase in RNAPII binding differs from that found in cells arrested in mid-S phase following hydroxyurea (HU) treatment [14]. In this later case, the prolong arrest, coupled with additional epigenetic changes, does allow ~40% increase in RNAPII binding to replicated DNA. Transcription buffering remains highly efficient also in this case, but requires additional compensation processes triggered by the DNA replication checkpoint.

### Transient depletion of TFs from replicated DNA

RNA polymerase is recruited to DNA by specific transcription factors (TFs) that bind to *cis* regulatory elements in promoter regions. We asked whether, similar to RNAPII, TF binding to DNA is also insensitive to DNA dosage. To examine that, we chose three factors of general regulatory roles that bind a large number of targets at distinct genomic regions: Reb1, Abf1 and Rap1 [18–20].

To define the binding dynamics of these TFs during S-phase, we released cells from G1 arrest, and followed their progression along S phase, sampling the culture at three-minute time intervals. Synchronized progression through S phase was verified using DNA staining (Figure 2A). TF binding profiles were measured by Chromatin Endonuclease Cleavage followed by Sequencing (ChEC-Seq) [21, 22]. Binding profiles were highly correlated between the different repeats and time points, and were consistent with previous data analyzing the binding pattern of the same factors in unsynchronized cultures (Figure 2B) [22]. Consistent with previous works, Reb1 and Abf1, were found preferentially ~130 bp upstream of the transcription start site (TSS), in regions that were largely depleted of nucleosomes, while Rap1 showed a more dispersed pattern along promoters (Figure 2C-E) [23]. Representative examples of the binding dynamics of the three TFs during the time course is shown in Figure 3A. Focusing on early replicating regions, we noted a transient depletion of TFs during early replication (Figure 3B). This reduced abundance lasted for 3-6 minutes for all three TFs and was more pronounced than that observed for RNAPII. Therefore, it appears that TFs are evicted from regions that are being replicated, presumably to prevent collisions with the DNA replication machinery, and then re-bind within minutes after the eviction.

**Figure 2:**
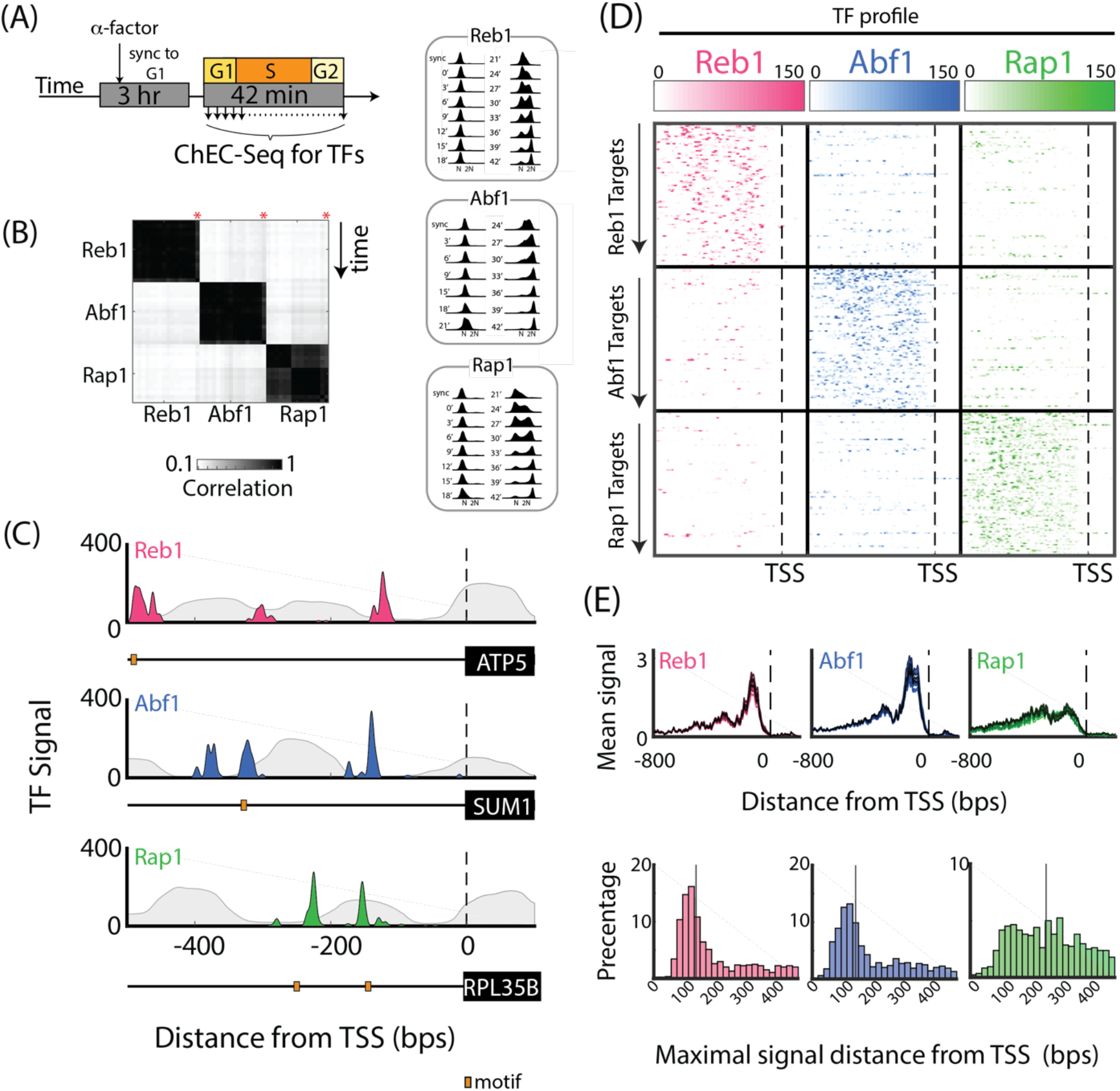
Genomic localization of the transcription factors Reb1, Abf1, and Rap1. (A) *Experimental scheme:* Yeast harboring MNase fused to the TFs Reb1, Abf1, or Rap1, were released from G1 arrest (left). The released cultures were sampled at 3-minute time intervals and processed for mapping TF binding pattern (Methods). Synchronized progression was verified using DNA staining (right). (B) *TF binding profiles are consistent between time points and repeats:* Shown are the Pearson correlations of the sum of signal on promoters, between all samples, time points and repeats. Correlation with previous datasets, corresponding to unsynchronized cultures [22] are also shown (red *) (See S2A-C). (C) *Representative binding profile to target genes:* Averaged binding intensities of each TF across all time points to the indicated gene promoters are shown. Location of sequence-predicted binding motif is indicated (orange box). Grey background depicts nucleosomes pattern in logarithmic growth as measured by MNase-Seq (see Methods; additional examples in S2E-G). (D) *Reb1, Abf1, and Rap1 bind to distinct sets of promoters:* The 100 promoters showing the strongest binding by each TF were selected. The binding patterns of the three TFs to the 300 selected promoters, aligned by their TSS (dashed line) are shown. Note the limited overlap between binding of the different factors. See Fig. S2D for all targets. The gene targets of each TFs are listed in supplementary table 1. (E) *Binding of Reb1, Abf1, and Rap1 along gene promoters:* Genes were aligned by their transcription start site (TSS). Shown is the average binding to all gene promoters (top), and the distribution of distances between the location of the maximal binding signal at each promoter to the TSS (bottom).

**Figure 3:**
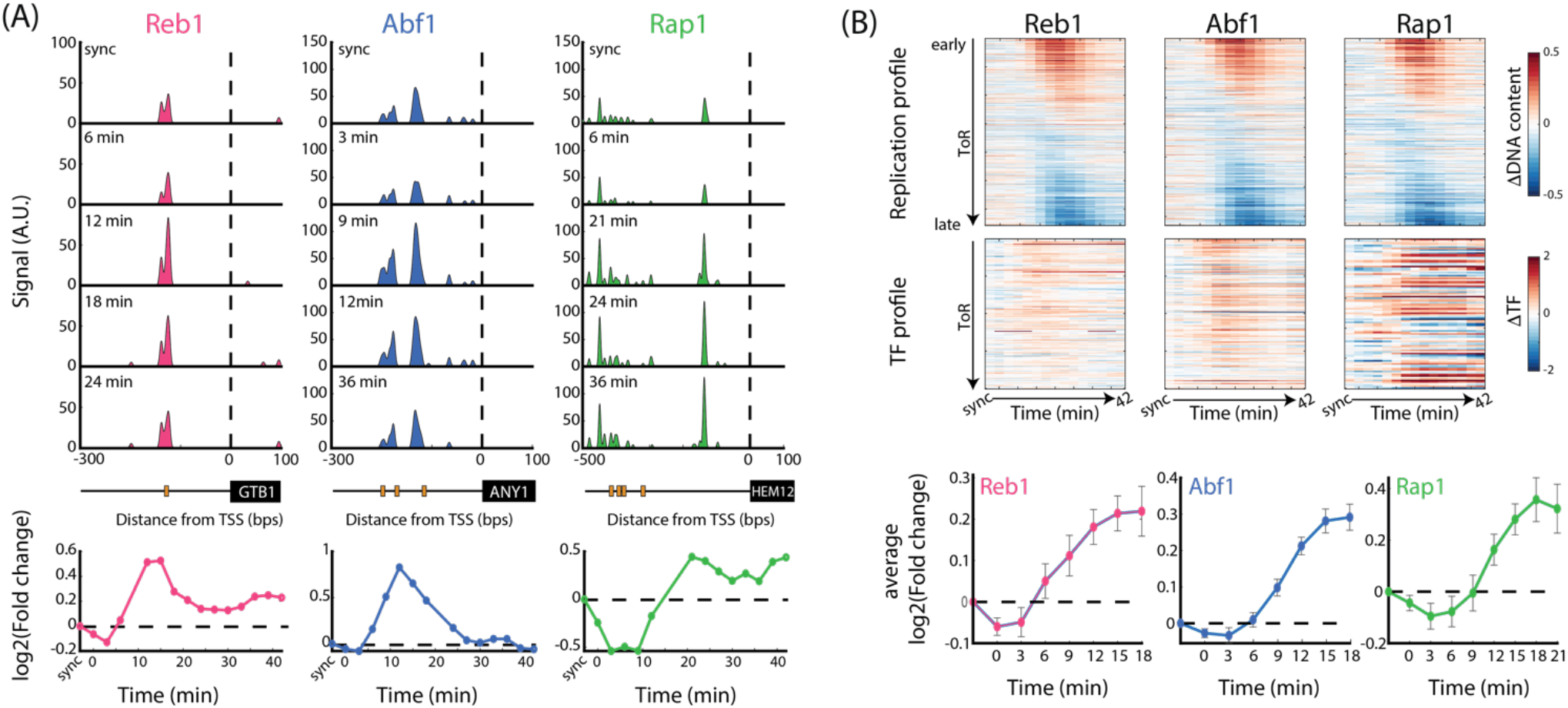
Transient eviction of TFs from replicated DNA. (A) *Temporal change in TF binding intensities at individual promoters:* A representative example of the binding dynamics of each TF, and its pattern along the promoter, is shown. The promoter was aligned according to the gene’s TSS, and the binding to the upstream sequence is shown (top, as in Figure 2C). Locations of binding motifs of the transcription factor is marked with orange boxes (middle panel). The signal over the of 500bps upstream of the TSS was summed, and the temporal change in this signal was calculated by normalizing by the synchronized time point (bottom). (B) *Temporal dynamics of all bound promoters:* Bound promoters (targets) were ordered according to their time of replication (ToR). The top panel shows the replication profile each gene, as calculated by measuring the (log2) fold-change in DNA signal on a 10kB region around each gene and compared to the synchronized time point (top). The top row is the replication profile of the earliest replicating target promoter. Note that since the replication profiles are measured using DNA sequencing, the late replicating promoters show a relative decrease in signal when early genes are replicated. The middle panel depicts the TF binding signal, as measured by ChEC-Seq over the 500bps upstream of the gene’s TSS, of the 200 earliest-replicating target genes for Reb1 and Abf1, and the 100 earliest-replicating target genes for Rap1. The average TF signal along time, and the standard error, over the same genes are presented in the lower panel.

### TF binding to replicated DNA increases in proportion to the increase in DNA dosage

Next, we examined whether, following their transient eviction, the binding of TFs to replicated promoters increases with the increasing DNA content. Using the ToR-based analysis described above (Figure 1D), we find that all three factors show biased binding to replicated DNA (Figure 4A). Thus, binding of Rap1 and Reb1 to replicated DNA increased in proportion to the increase in DNA content, while the increase in DNA binding of Abf1 surpassed the increase in DNA dosage by ~50%. The post-replication chromatin environment is therefore permissive for TF binding and may in fact become more accessible for binding of certain factors, such as Abf1. When repeating the analysis for RNAPII binding, focusing on the gene targets of each TF (Figure S3F), we find similar dynamics to those observed in our global analysis (Figure 1E), and thus, RNAPII binding to replicated genes is inhibited.

**Figure 4:**
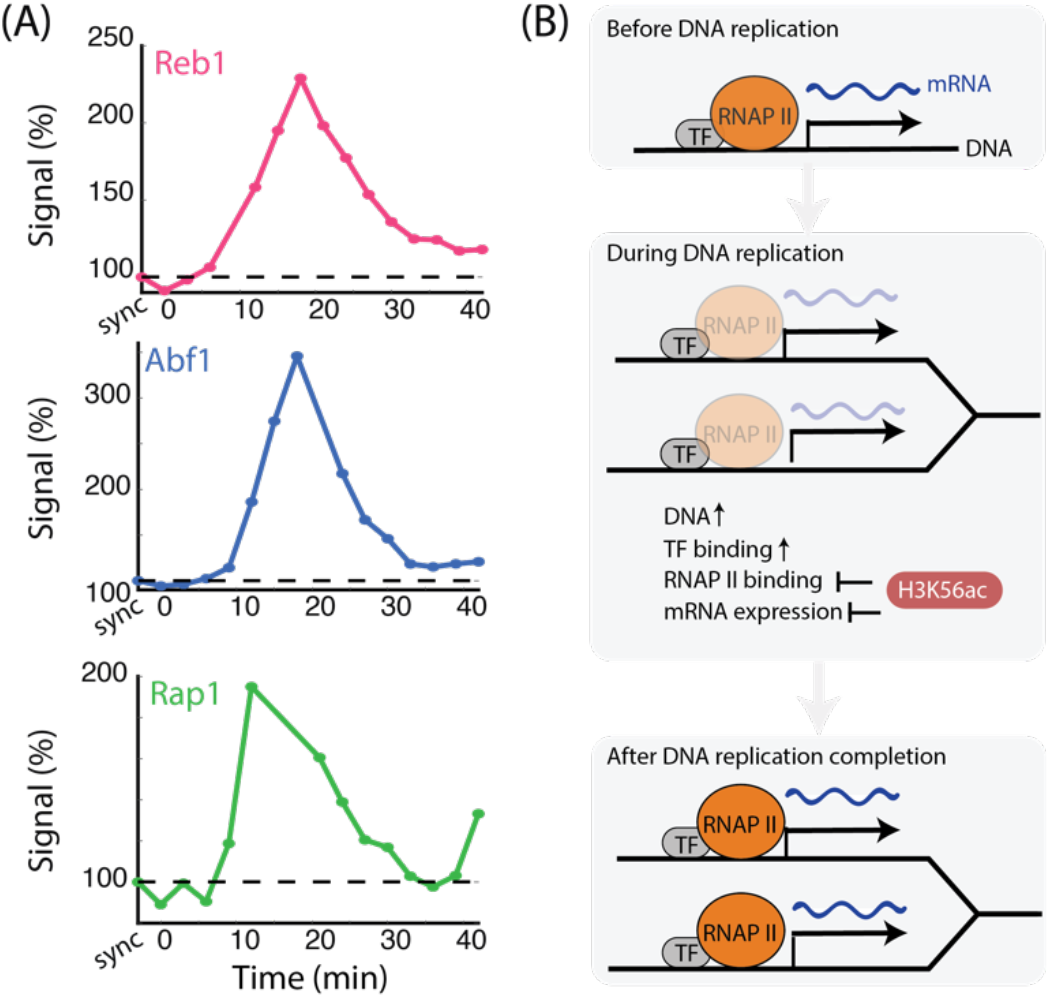
TF binding to replicated DNA increases with the increasing DNA content. (A) *Increasing TF binding intensity with increasing DNA dosage:* The binding profile for each transcription factor was calculated over its corresponding set of target genes. The binding was summed on the promoter region, defined as 500 bps upstream of the TSS. The dependency of TF binding on ToR along time was calculated for each time point, as in Figure 1E. See also Figure S3 for TF binding in rtt109-deleted cells. Analysis of Pol2 binding over the same gene set is shown in Figure S3F. (B) *Model:* Prior to DNA replication, TFs bind their target promoters, recruit RNAPII, and subsequently mRNA is transcribed. During DNA replication, TFs are temporarily evicted from the DNA template, but then re-bind rapidly to the replicated DNA, poised to activate transcription. Despite TFs binding the replicated promoters, RNAPII is not recruited to replicated genes, resulting in buffering of the imbalance in gene dosage that occurs during S-phase.

The increased binding of TFs to replicated regions contrasts the suppression of RNAPII binding and gene transcription from these regions. Still, we asked whether this binding increases even more in mutants that are deficient in transcription buffering. To test that, we examined the S-phase binding patterns of Abf1 and Reb1 in cells deleted of the histone acetyltransferase *RTT109*. Both factors showed the precise same binding dynamics as observed in wild-type cells (Figure S3D&E). Thus, H3K56ac, while suppressing transcription from replicated DNA, does not affect TF binding.

## Discussion

Differential binding of regulatory proteins to DNA is critical for the ability of a single genome to dictate different phenotypes. DNA replication challenges this organization in at least three ways. First, the passing of the replication fork requires the removal of obstacles that may impede fork progression. Second, new histones, introduced for wrapping the newly synthesized DNA, carry a unique set of marks, which could influence regulatory factor binding. Finally, the changes in the relative abundance of different genomic regions, introduced by differences in their replication timing, may draw regulatory proteins preferentially to early replicating regions at the expense of late replicating ones. We recently showed that these biases have little effect on gene expression, both during normal S phase and in cells arrested with their genome partially replicated [13, 14]. Here, we examined how these biases affect proteins that bind DNA to regulate transcription: RNAPII, and specific transcription factors.

Previous studies indicated that during replication stress, such as HU-treatment, RNAPII is evicted from DNA for ~60 minutes [24]. Our genome-wide analysis of RNAPII binding during normal S phase revealed a highly stable pattern, with little, if any, detectable depletion upon the passing of the replication fork. Our time resolution (3 minutes) cannot resolve whether RNAPII is transiently evicted and rebinds rapidly, or whether it remains in the close vicinity of the replication fork. In either case, the passing of the replication fork does not significantly perturb the overall RNAPII binding pattern, neither in the promoters, where it awaits initiation signals, nor within the coding regions, where it is actively elongating to produce the mature transcript.

Previous studies also proposed that specific TFs are evicted from DNA during replication. This was based on the pattern of protected DNA-fragments [7], or the transient inaccessibility of chromatin during DNA replication as recently measured by ATAC-Seq [25]. In addition, by motif analysis, TFs have been correlated with Okazaki fragment processing, implying that TFs may bind replicated DNA rapidly [26]. However, neither the identity of evicted factors, nor the dynamics of this eviction, were described. Our data, examining three specific TFs with a wide spectrum of targets, did indicate such an eviction, showing a depletion of these factors from early replicated regions that was more pronounced than that observed for RNAPII. This depletion, however, is transient, lasting only several minutes.

Following the passing of the replication fork, DNA dosage increases, and it is present within a unique chromatin environment, characterized by a distinct pattern of histone marks. We asked whether replicated DNA remains accessible for binding regulatory factors. Previous studies suggested that the unique chromatin environment characterizing replicated DNA is less permissive for RNAPII binding, based on the observation that replicated promoters are transiently occupied by nucleosomes [6–8]. Consistent with that, we find that RNAPII remains largely insensitive to the increase in DNA content, showing only a limited increase in binding to replicated regions. However, the kinetics by which promoters regain the correct nucleosome positioning appears too rapid to explain the persistent suppression of RNAPII binding we observe. Further, TFs binding to replicated promoters does increase with the increasing DNA content. Therefore, the replication-dependent chromatin environment remains permissive for binding of TFs. Of note, for at least one of these factors, binding to replicated regions surpassed the increase in DNA content, suggesting that it binds replicated DNA with increased efficiency compared to non-replicated DNA. This excessive binding of TFs to replicated promoters may explain the transcriptional spike upon mitotic exit that was recently observed in human cells [27].

Taken together, our work provides new insight into the dynamics of RNAPII and of specific TFs during DNA replication. While TFs are evicted from the DNA during replication, they regain binding rapidly and are poised on replicated promoters to initiate transcription in the daughter cells. The unique chromatin environment during replication, however, limits the ability of these factors to recruit RNAPII to the replicated genes, explaining the observed buffering of the gene dosage imbalance at the level of mRNA expression (Figure 4B).

## Materials & Methods

### Strains and Plasmids

For ChEC-seq experiments in wild-type *Saccharomyces cerevisiae*, a BY4741 was tagged with MNase by amplifying a cassette from pGZ108 [22]. Yeast were transformed with a PCR fragment amplified with the following primers – For Reb1 - 5’-TGATTATTTTAGCTCCAATATTTCAATGAAAACAGAAAATggtcgacggatccccgggtt and 5’-TTATTGAGTTTTTCGCTTTCACCAATTATATTTTCCGGAAtcgatgaattcgagctcgtt. For Abf1 – 5-CCTTTCTGATGAAAACATTCAACCAGAATTAAGAGGTCAAggtcgacggatccccgggtt 5-AGAAACATGAGAAAAATAGCTC GTCTTCTCAACTGGGTATtcgatgaattcgagctcgtt. For Rap1–5-AATGGAAATGAGGAAAAGATTTTTTGAGAAGGACCTGTTAggtcgacggatccccgggtt and 5-GT AAAAT AAGTT AAACAAT GAT GTT ACTT AATT CAATT ACtcgatgaattcgagctcgtt and selection on plates with G418. Reb1-MNase and Abf1-Mnase strains were further transformed with a PCR fragment amplified from pYM24 [29] to delete *RTT109* (Figure S3) replacing the gene’s ORF by transformation and growth on plates containing G418 and Hygromycin B. For MNase-seq experiments the wild-type *Saccharomyces cerevisiae*, BY4741, was used.

### ChEC-Seq during the cell cycle

Cell cycle synchronization using α-factor was done as previously described [5]. Briefly, cells were grown in YPD overnight at 30°C and inoculated in fresh medium to OD600 of 0.05. When reaching an OD600 of 0.12, cells were washed from the media by centrifugation (3000 rpm, 5 min). Cells were then resuspended in an equal volume of fresh warm YPD with α-factor to a final concentration of 5 μg/mL. Next, the yeast culture was divided into 16 (experiment set #1, Figure 4A, S3D-E) or 4 (experiment set #2, Figure S3B-C) separate 50-mL tubes with a ventilated cap (CELLSTAR CELLreactor filter top tube, Greiner Bio-One, 227245), each containing 35 mL of yeast culture. Each tube contained the material used for ChEC-Seq in a single time-point and DNA staining for flow cytometry. Cells were incubated for 3 hours at 30°C with α-factor in an incubator and then transferred to a water bath orbital shaker 30 min before the end of synchronization (MRC, WBT-450). Every 3 min for 42 min, one tube was taken out of the bath orbital shaker and washed twice from α-factor by centrifugation (4000 rpm, 1 min) and resuspension in fresh, warm YPD. Following two washes, cells were resuspended in an equal volume of fresh, warm YPD and returned to the bath shaker to grow at 30°C. The first sample returned to the bath shaker is the last sample in the time-course (first released–last time-point in time-course). The second-to-last sample in the time-course was released and immediately processed, termed the “0 minutes” sample. The last sample in the time-course was not released from α-factor and was termed “synchronized.” Following release from synchronization, 0.5 mL of each sample was aliquoted to a different tube to be used for DNA staining and flow cytometry. Samples for DNA staining were centrifuged for 10 seconds in 13000 rpm, sup was discarded, and pellet was resuspended and fixated with 70% ethanol. The remaining culture was used for CheEC-Seq, as described previously (Zentner et al., 2015), with minor modifications. Briefly, cells were pelleted at 1500g, transferred to a deep-well 96-well plate, and then washed three times with 1 ml Buffer A (15 mM Tris pH 7.5, 80 mM KCl, 0.1 mM EGTA, 0.2 mM spermine, 0.5 mM spermidine, 1 × Roche cOmplete EDTA-free mini protease inhibitors, 1mM PMSF). Cells were then resuspended in 200μl Buffer A containing 0.1% digitonin and permeabilized at 30 °C for 5 min. CaCl2 was added to a final concentration of 2 mM for 30 seconds at 30°C. Next, stop buffer (400 mM NaCl, 20 mM EDTA, 4 mM EGTA) and 1% SDS was added and vortexed. Proteinase K (100μg, Sigma-Aldrich) was then added and incubated at 55°C for 30 min. Subsequent nucleic acids isolation, RNAse A treatment, and subsequent DNA clean-ups were done as previously described (Zentner et al., 2015). For the second set of experiments (Figure S4), small adjustments were made in the library preparation in order to increase yield: Reverse SPRI clean-up for enrichment of small DNA fragments was done (0.8X). Ethanol precipitation was performed during the clean-up steps instead of using S300 spin columns (120uL EtOH 96% and 5uL of Sodium acetate (3M) were added to ~50 uL of sample, vortexed and precipitated at −80C for > hour), followed by a final SPRI cleanup (1X). ChEC libraries were indexed [30], pooled and sequenced on Illumina NextSeq500 for single 50bps reads.

### MNase-Seq

For profiles of nucleosome occupancy in logarithmic growth, MNase-seq was performed as previously described [31]. Cells were grown over night, then diluted and grown for 5-6 hours shaking at 30°C to reach OD=0.5. 10ml of cells were fixated for 15 min in 1% formaldehyde shaking in RT. Cell pellets were washed, and treated with zymolyase (Amsbio, 120493-1) for 25 minutes in 30°C to generate spheroplasts. Then, spheroplasts were subjected to MNase digestion (Worthington, LS004797) for 20 minutes in 37°C. MNase treatment was stopped using a stop buffer (220mM NaCl, 0.2% SDS, 0.2% sodium deoxycholate, 10mM EDTA, 2% Triton X-100). Cells were reverse cross linked: RNase (Sigma, R4875) treatment in 37°C for 30 min and then Proteinase K (Sigma P2308) in 37°C for 2 hours. Samples were incubated overnight at 65°C and DNA was purified using SPRI beads with a ratio of 2X. DNA libraries were indexed [32] pooled and pair-end sequenced on Illumina NextSeq500.

### Flow cytometry

To verify cell-cycle synchronization efficiency and position along the cell cycle, we performed DNA staining of samples from every time-point using flow cytometry. Briefly, cells were washed twice with 50 mM Tris-HCl (pH=8), resuspended in RNase A for 40 min in 37°C, washed twice with 50 mM Tris-HCl (pH=8), and resuspended in Proteinase K for 1 h incubation at 37°C. Then, cells were washed twice again, resuspended in SYBR green (S9430, Sigma-Aldrich; 1:1000), and incubated in the dark at room temperature for 1 h. Then, cells were washed from the stain and resuspended in 50 mM Tris-HCl (pH 8) and sonicated in the Diagenode Bioruptor for three cycles of 10 sec on and 20 sec off in low intensity. Finally, cells were taken to FACS for analysis using the BD LSRII system (BD Biosciences).

### Gene expression data and analysis

Gene expression data and mRNA synthesis rates for wild type yeast (BY4741) and *Δrtt109* measured along the cell-cycle following release from a-factor synchronization previously published by our lab was used [13]. Note, the experimental scheme and time points are the same as in the current study. Briefly, cells were released, after washing, from 3 hours in YPD with 5μg/mL a-factor into fresh YPD and samples for RNA-Seq and flow-cytometry were harvested, every 3 min for 39 min, and then every 6 minutes for a total of 135 min. Expression levels at each time-point were divided by the expression in the synchronized time-point, log2-transformed, and then averaged on the group of genes or single genes.

### Pol II ChIP-Seq and genomic DNA data

Pol II and genomic DNA data along S-phase from a previously published dataset from our lab was used (Bar-Ziv et al., 2016). Briefly, a wild type yeast culture (W303) progressing synchronously following a-factor synchronization, was used. Cells were grown in 24°C, arrested by α-factor for 2.5 hours, and shifted to 34°C for half an hour before release from G1 arrest. At each time-point, 35 mL of yeast were used for ChIP-seq, with chromatin from 10 mL of culture used per antibody. Tubes were taken out of the bath shaker and fixated at the same time, as each sample has been released at a different time, in 3-min intervals. The antibody used to probe RNA Pol II is Pol II (CTD) 8WG16 (Covance, MMS-126R). The cells were fixated and processed further as previously described. Samples were harvested from the synchronized culture, and then from 6 min after release taken every 3 min for 45 min total after release from synchronization. Reads were aligned using Bowtie (parameters: –best –m 1) to a combined *S. cerevisiae* and *S. pombe* genome. This experimental scheme, that used spiked-in *S. pombe*, was not used in other experiments presented here. The spike-in method was an experimental method that was used to probe total levels of chromatin marks in our previous study, but was not needed for calculations in the current study (see note in Bar-Ziv et al., 2016a) *Genomic* tracks were calculated by extending the aligned reads to cover 200 bp and adding +1 to each covered location. All tracks were normalized to have a total signal of 1,000,000. For analysis of RNAPII binding by ToR bias, regulated genes were taken out of the analysis (as in Voichek et al., 2016). Genes that are known to be transcriptionally regulated under the conditions used in our experimental system, defined here as “regulated genes”, were excluded from the analysis. This group of genes includes cell cycle genes (“cell-cycle (G1)”, “cell cycle(G2/M)” and “cell cycle(CDC15)”), genes regulated in response to mating-factor synchronization (“mating”) and genes responding to stressed condition (“protein synthesis”, “stress”,”rRNA processing”, “ESR induce” and “ESR reduce”) [33, 34]. All genes are listed in Table S1.

### ChEC-Seq processing and analysis

Reads were aligned using Bowtie2 to a *S. cerevisiae* (reference genome, cerR64). ChEC-Seq tracks representing the enrichment of every locus in the yeast genome were calculated for all samples. Genomic tracks were calculated by adding +1 to each genomic location corresponding to the first nucleotide in each read and normalized to have a total signal of 10,000,000 to control for sequencing depth. The signal on each promoter (500bps upstream of the TSS, [28]) was summed. Targets genes were defined by genes with a positive cumulative signal in their promoter above noise. For temporal dynamics, for each time point the total amount of signal was normalized, and the log2 fold change between each time point and the synchronized time point was calculated. Genes that were identified as gene targets of the TF reproduced previous results of gene targets for each TF as in previously published datasets (Figure S2). Several samples for experiment set #1 were discarded from further analysis due to technical issues, such as low alignment rate: Reb1-Mnase 9 min, Abf1 21-Mnase min, Rap1-Mnase 15&18 min, Reb1-Mnase Δrtt109 15&18&30 min, Abf1-Mnase Δrtt109 9&12&18 min. For analysis of TF binding by ToR bias, regulated genes were taken out of the analysis (as in Voichek et al., 2016).

### Metagene profiles

For RNA Pol II binding, metagene analysis was done as follows: Taking the signal 500 bps upstream of the transcription start site (TSS) for every gene in this profile [28]. The signal found between the TSS and the transcription termination site (TTS) was binned into 50 equal-sized bins to be able to compare genes of different lengths. Signal for all genes was then separated according to expression levels, and averaged to get the average pattern. For TF binding, metagene analysis was done as follows: taking the signal 800 bp upstream of the transcription start site (TSS) for every gene in this profile, and 100 bps downstream of the TSS. Signal for all genes was averaged to get the average pattern.

### Motif analysis

For finding the motifs of TFs on the presented promoters, the top-scoring motif from The Yeast Transcription Factor Specificity Compendium [35] was searched along the gene’s promoter using FIMO [36].

### Quantifying signal along DNA replication

To quantify the change in signal along DNA replication (as shown in Figure 1D-G, Figure S1C), the log2 fold change of each gene (for RNAPII, DNA replication, gene expression, and mRNA synthesis), compared to the synchronized time point, was plotted. The data was then fitted using first-degree polynomial fitting. The slope was then linearly transformed, so that the maximum increase in genomic DNA during DNA replication would be 2. For RNAPII, the signal on the promoter region (500 bps upstream of TSS), or the signal on the ORF was summed. For DNA content, the region of 10kB around the TSS of the gene was summed. For TFs, the same calculation was done, calculated only for target genes. For TFs, the binding was summed on the promoter, 500 bps upstream of the TSS. Note that “regulated genes” (see above, Table S1) were discarded from the quantification.

### Time of replication (ToR) and gene groups

Replication timing data of DNA from Yabuki et al., 2002 was used to define gene replication time by assigning each gene the replication time closest to its 5’ end. Gene groups for RNA analysis (Figure 1) G2/M genes and G1 genes were taken from Ihmels et al., 2004. For the histone genes group, all 8 histone genes were used.

## Availability of data and materials

The ChEC-seq and MNase-Seq dataset supporting the conclusions of this article are available in the NCBI Sequence Read Archives repository, accession number BioProject PRJNA542378.

## The authors declare no competing interests

### Author contributions

R.B.-Z. and N.B. conceived the study and designed experiments; R.B.-Z, S.B., and M.C. performed experiments; R.B.-Z and S.B., analyzed the data; R.B.-Z, S.B., and N.B. wrote the manuscript; N.B. supervised the research.

## Acknowledgments

We thank O. Lupo, Y. Gordon, Y. Voichek, and M. Carmi for technical assistance. We also thank members of our laboratory for fruitful discussions and comments on the project. This work was supported by the ISF and Minerva.

