## Supplemental Figures for "Transcription-factor binding to replicated DNA"

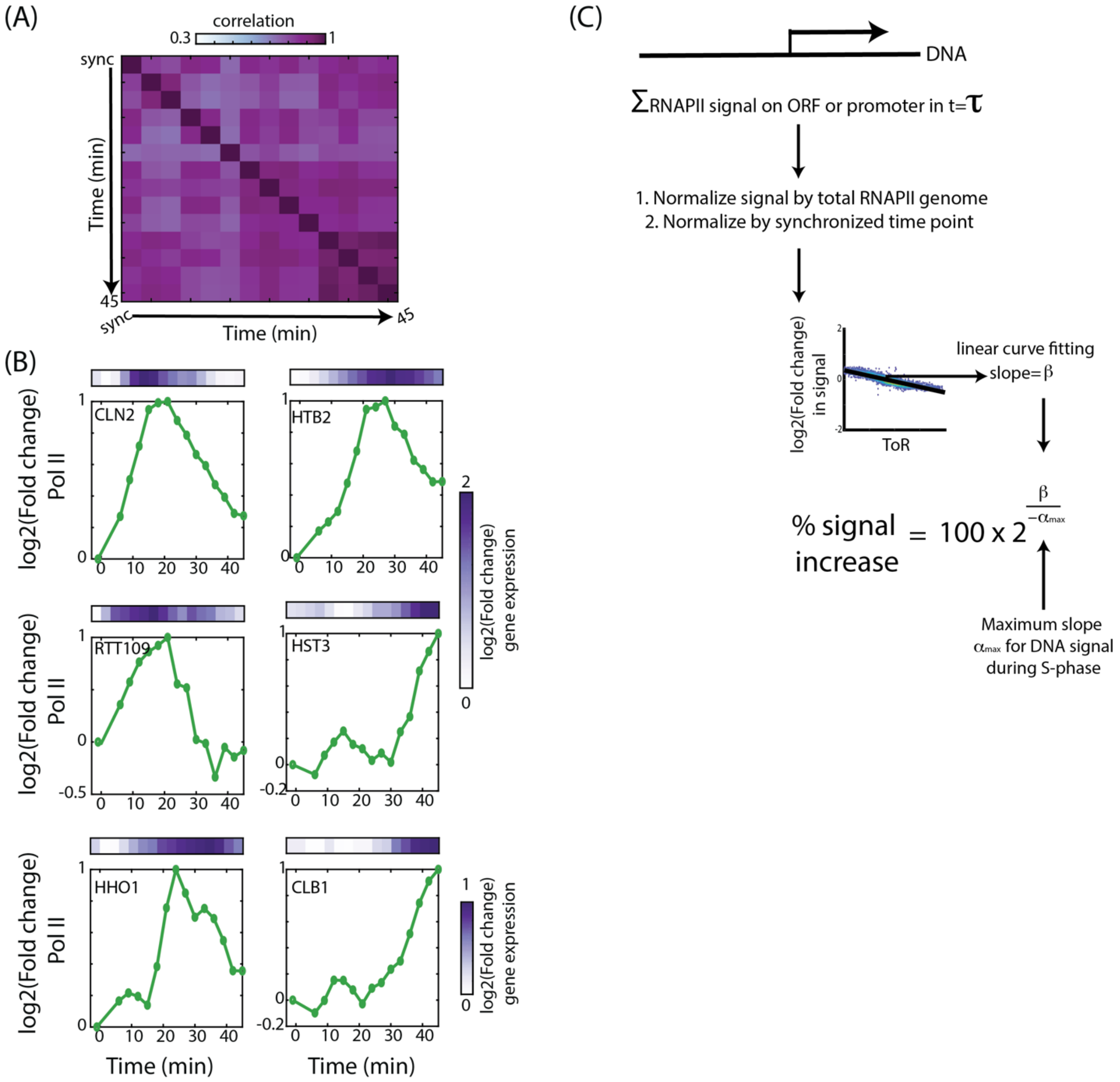


**Figure S1:** ***The recruitment of RNA polymerase II to replicated DNA.***

(A) *Correlations between data points:* The Pearson correlation between time points of samples of RNAPII ChIP-Seq.

(B) *RNAPII binding to regulated genes:* Examples of the change in Pol2 binding to regulated genes along the time course shows the expected cell-cycle dynamics; G1/Late-G1 genes (*CLN2* and *RTT109*), S-phase histones (*HHO1* and *HTB1)*, and G2/M genes (*HST3* and *CLB1)*. The average (log2) fold change in binding dynamics of RNAPII on the indicated gene (bottom), and the log2 fold change in mRNA levels of the gene (heatmap, top), upon release from $\alpha$-factor synchronization, along the time course, normalized to the synchronized time point.

(C) Calculating the dependency of RNAPII on time of replication: For each time point $\tau$, the RNAPII signal on each ORF or promoter was summed, and normalized by the total signal of RNAPII on the genome. Then, the signal was normalized to the synchronized time point, to get the fold-change increase in signal per gene. Next, the fold change in RNAPII signal on all examined genes was plotted against their ToR, and by linear curve fitting the slope (per time point $\tau$ along the time course) was extracted. Finally, the slope of the curve was divided by the negative of the maximum slope (generated similarly for DNA content), and converted to percentages so that the synchronized time point is a 100%.

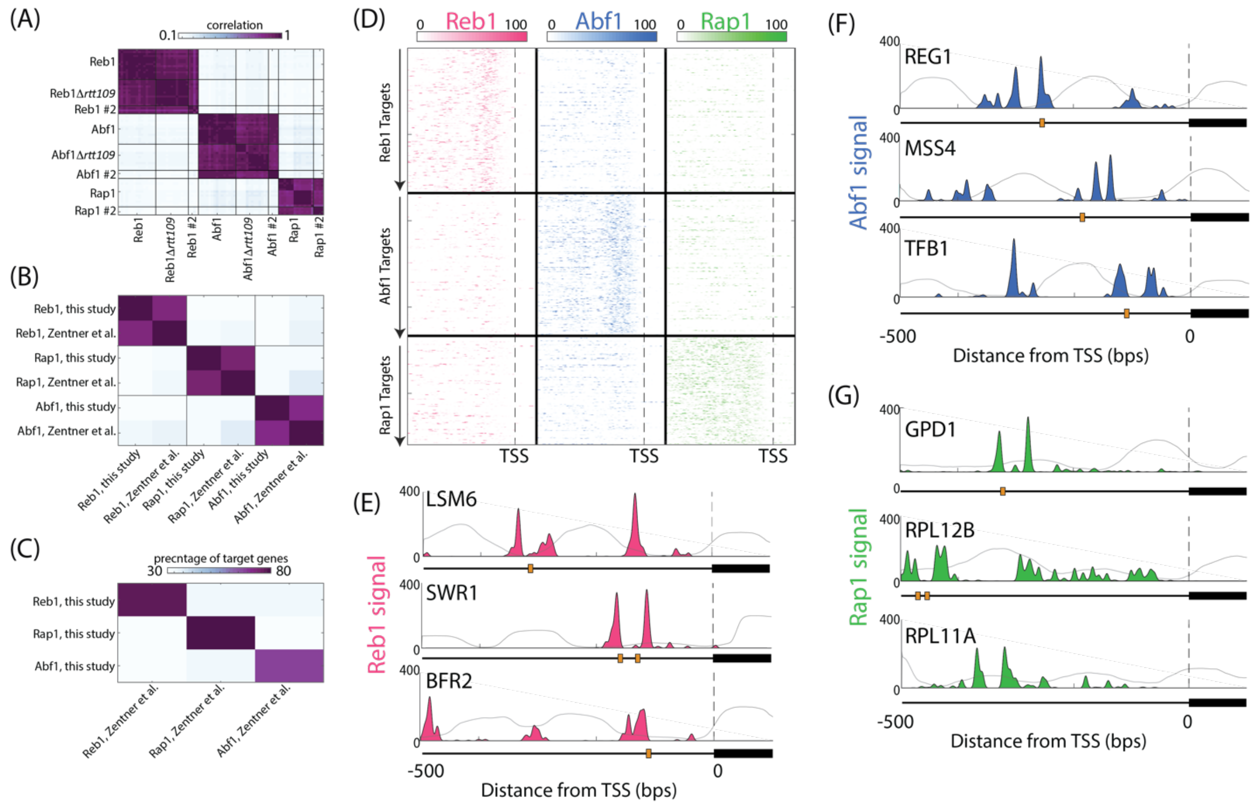


**Figure S2:** ***Measuring the binding of transcription factors during DNA replication***

(A) *Correlations between experiments:* The signal on each promoter for each TF, in each time point, was summed and cross-correlated to all other samples.

(B) *Correlations with previous data* (Zentner et al., 2015)*:* Median sum on promoter of each TF was calculated, and then correlated to the binding data produced by Zentner et al. for yeast probed during logarithmic growth.

(C) The percentage of overlap in identified target genes for each TF in this study vs Zentner et al, and between the different TFs, is shown.

(D) *Spatial binding patterns of TFs:* As Figure 2D, the spatial binding pattern of the indicated TFs is shown, for all identified targets of each TF. Patterns are shown in distance from the transcription start site (TSS).

(E) *Single-gene examples:* As Figure 2C, the binding patterns of three target genes of each transcription factor are shown. The location of motifs in the promoter (orange box) and the transcript (black box) are indicated. Grey background signal represents the pattern of nucleosomes in logarithmic growth.

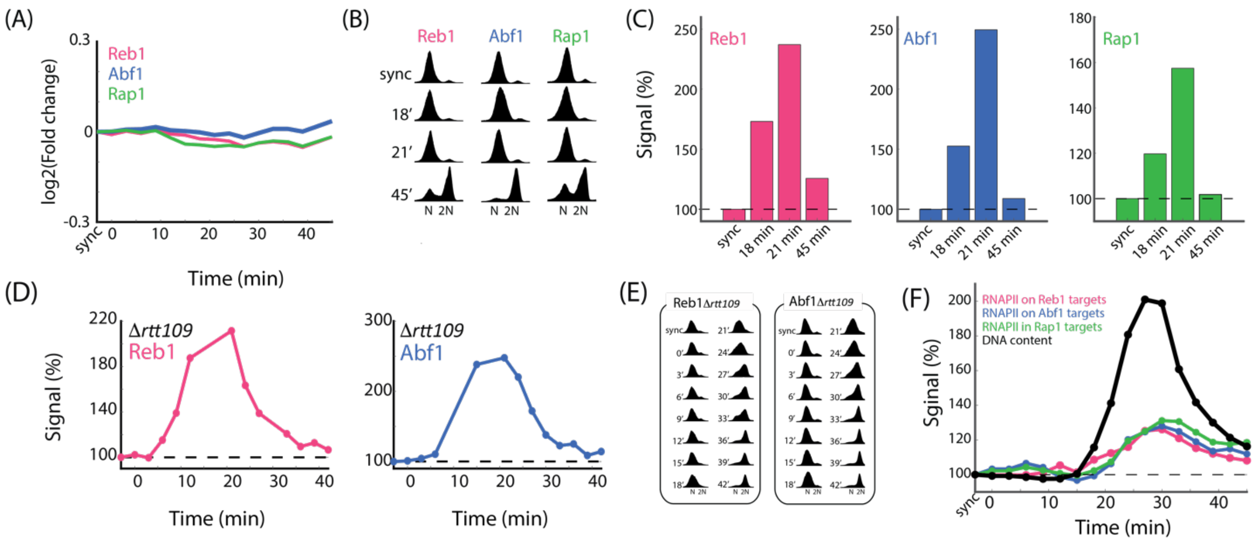


**Figure S3:** ***TFs re-bind to replicated DNA***

(A) *Little fluctuations in gene expression of TF targets during S-phase*: The log2 fold change in expression levels in a wild type strain, as measured by RNA-seq, of the gene targets of each TF is shown. Data from Voichek et al., 2016.

(B) *Cell cycle progression:* Synchronized progression for results shown in Figure S4C. Cell cycle progression was verified by total DNA content profile using fluorescence-activated cell sorting (FACS).

(C) *TFs binding to replicated DNA is not buffered:* The binding dynamics of each TF on all its targets was calculated as in Figure 1E (biological repeat #2).

(D) *Increasing TF binding intensity with increasing DNA dosage also when deleting Rtt109:* Same as Figure 1E for the indicated binding profiles. Synchronized progression was verified using DNA staining (E).

(F) Minor sensitivity of *RNAPII to gene dosage when examined only on gene targets of each transcription factor:* As in Figure 1E, shown are the dependencies of RNAPII binding to ORFs on ToR along time. The analysis was done for the set of target-genes of each transcription factor.
